# Geological drivers of diversification in Papuan microhylid frogs

**DOI:** 10.1101/2022.09.15.508064

**Authors:** Ethan C. Hill, Diana F. Gao, Dan A. Polhemus, Claire J. Fraser, Bulisa Iova, Allen Allison, Marguerite A. Butler

## Abstract

Studies of the Papuan region have provided fundamental insights into both the evolutionary processes generating its exceptional biodiversity, as well support for alternative hypotheses of geological history. Lying at the junction of five tectonic plates, this region has experienced a turbulent geological history that has not only produced towering mountains allowing elevational specialization, and island archipelagos of varying distance promoting vicariance, but also active margins where land masses have collided and been subsequently rifted apart creating a mosaic of intermixed terranes with vastly different geological histories which may influence the evolutionary history of its biota. Asterophryine frogs are a hyperdiverse clade representing half the world’s microhylid diversity (over 360 species) centered on New Guinea and its satellite islands. We show that vicariance facilitated by geological history, and not elevational specialization best explain this far and wide distribution of a clade that should have poor dispersal abilities. Thus, some of the predictions of island biogeography theory are supported if informed by geological history. We recovered a mainland tectonic unit, the East Papua Composite Terrane (EPCT), as the center of origin for Asterophryinae and no fewer than 71 instances of what appear to be long-distance dispersal events, 29 of which are between mainland regions, with 42 from the mainland to the islands, some presently as far as 200 km away from source populations over open ocean. Furthermore, we find strong support for a “Slow and Steady” hypothesis for the formation of the northern margin of New Guinea by many separate accretion events during the Miocene, over other major geological alternatives, consistent with the 20 M year age of the clade and arrival via the EPCT. In addition, the historical biogeography of our frogs strongly support an affiliation of the Louisiade Archipelago and Woodlark Island with the Owen Stanley Mountain range on the EPCT, and the recent proximity of the large New Britain island. Our results show that Asterophryinae did not have to repeatedly and independently disperse across and large ocean barriers to the offshore islands, but that the current distribution can be explained through vicariance and short-distance oceanic dispersal as historical land connections disappeared and islands slowly became separated from each other. We show that islands have a life history, undergoing changes in area through island-building and erosion, but also change in distance from other land masses, with consequent opportunities for dispersal, isolation, and cladogenesis of their biotas. More broadly, we can begin to see how the geological history of the Papuan region can result in the rapid accumulation and staggering number of extant species.

---

> *“The distributions of organisms not adapted for long-distance dispersal are good evidence of past land connections*.*”*
>
> — Biogeographical principles advocated by Sir Alfred Russell Wallace
>
> Source: Brown and Lomolino (1998)

Wallace (1876, 1880) proposed that to understand the fascinating distributional patterns in the South Pacific requires a consideration of both natural selection and earth history (Hazzi et al., 2018; Pellissier et al., 2018). The island of New Guinea, lying in an intersection zone between Australian and Asian faunal and tectonic provinces, contains among the highest degrees of endemism of any place on earth (Brooks et al., 2006; Gressitt, 1982) and provides a case in point. This large island was formed over the past 30 MY (Polhemus, 2007; Baldwin et al., 2012), when many taxa dispersed to New Guinea and diversified via founder event speciation, for example: ants (Toussaint et al., 2015), butterflies (Toussaint and Balke, 2016), damselflies (Kalkman et al., 2018), water bugs (Polhemus and Polhemus, 2004), leaf insects (Bank et al., 2021), birds (Jonsson et al., 2017), lizards (Tallowin et al., 2018), and plants(Segar et al., 2017). The largest tropical island in the world, its mountain peaks rise as high as 4800m to provide a broad range of climate and habitat variation set amid a rugged topology, which has driven local adaptation (Allison, 2009). Its elevational gradients are famously associated with species diversity across diverse taxa (Tallowin et al., 2017) including: birds (Diamond, 1973), plants (Hoover et al., 2017), insects (Beck et al., 2017; Toussaint et al., 2014; Souto-Vilaros et al., 2020; Cozzarolo et al., 2019), lizards (Slavenko et al., 2020), and frogs (Oliver et al., 2017). The surrounding region also contains numerous proximal island archipelagos, offering the perfect balance of isolation, adaptation, and migration potential for species to diversify. Indeed, early studies of the Melanesian ant fauna inspired the taxon cycle hypothesis (Wilson, 1959, 1961; Toussaint et al., 2015), as well as contributed to the theory of island biogeography (MacArthur and Wilson, 1967; Losos and Ricklefs, 2009; Mayr and Diamond, 2001), relating dispersal potential to island distance and speciation and extinction rates to island size.

Wallace’s second proposal, that a changing geology may shape the distribution and diversity of life, has received less study, but is supported by correlation between geological events and areas of high species richness – the rise of central high mountains is associated with vicariance in ants (Li and Li, 2018) and freshwater turtles (Georges et al., 2014), and signatures of presumed island fragmentation and accretion sequences are reflected in patterns of endemism in island insects (Boer and Duffels, 1996; Polhemus, 1996; Polhemus and Polhemus, 1998). These processes are amenable for testing in the greater New Guinea region.

Recent advances in phylogenetic methods incorporating timing informed by new geological data for New Guinea (reviewed in: Baldwin et al., 2012) have inspired testing of more explicit spatio-temporal hypotheses of distribution and diversity (e.g., Jonsson et al., 2011; Georges et al., 2014; Toussaint et al., 2014). For example, Toussaint et al. (2014) found support for recent orogeny driving the diversification of montane arthropods. Specifically, this dataset supported the geological model of Hall (1998, 2002) which predicts that as late as (*∼*5Ma), most of New Guinea was under water with only the central high mountains above sea level and available for colonization by terrestrial taxa. This model posits that collision between the Pacific and Australian plates resulted in uplift of the central cordillera and the rest of mainland New Guinea. While this “recent emergence” model works well to explain diversity patterns of young taxa centered on the central cordillera, there are additional geological hypotheses to consider, which can be tested with clades that are older and more widely distributed across New Guinea (but see: Cozzarolo et al., 2019).

New Guinea lies in one of the most tectonically active regions of the world, and while its geological history is complex, there is now general agreement that collision of plates and the accretion of island arcs played a significant role in building the island (reviewed in: Baldwin et al., 2012). Starting along the northern edge of the Australian Craton, the island formed over the Cenozoic with the additions of the East Papua Composite Terrane, the Fold Belt, the Accreted Terranes, and the Vogelkop Peninsula (Baldwin et al., 2012; Crowhurst et al., 1996; Davies et al., 1996, 1997; Dow, 1977; Pigram and Davies, 1987; Pigram and Symond, 1991; Hall, 2002; Holm et al., 2019; Quarles van Ufford and Cloos, 2005; Webb et al., 2014). These processes were initiated by the collision of the northward-moving Australian Plate with the west-northwest-moving Pacific Plate, with the additional interactions of the smaller Philippine, Caroline, and Solomon Arc plates (Hall, 1998; Klootwijk et al., 2003; Kroenke et al., 1984; Kroenke, 1996), and various other microplates that were shattered off along their edges during convergence. This resulted in sequential accretion of island arcs and subduction or obduction of oceanic and continental crust (Polhemus, 2007; Baldwin et al., 2012), to form an island of large size and complex topography, as well as a specific temporal ordering of land connections. The last major addition was the rotation and subduction of the South Bismarck Plate onto the northern margin as part of the Accreted Terranes, adding the Adelbert-Finnisterre terrane containing the Huon peninsula within the last several million years, and bringing the New Britain island arc into the closest proximity to the mainland that it has ever been (Baldwin et al., 2012).

Two additional geological hypotheses of particular relevance for biotic diversification, differ from the “recent emergence” hypothesis and each other with regard to the timing and assembly of the northern coastal terranes. The “Mobile belt” hypothesis posits that the Accreted Terranes were assembled offshore into a single unit during the late Oligocene (25-23 Ma), and subsequently docked onto the growing mainland in the mid-Miocene (15-11 Ma; Dow, 1972, 1977; Davies, 2012) giving rapid rise to the central high mountains in the Fold Belt (Quarles van Ufford and Cloos, 2005; Webb et al., 2014; Holm et al., 2019). Alternatively, several authors hypothesize that accretion along the northern coast has been a “slow and steady” process extending from the late Cretaceous to the Pleistocene (Pigram and Davies, 1987; Davies et al., 1996; Quarles van Ufford and Cloos, 2005). Under this scenario, the ECPT is a composite of terranes which accreted onto a displaced sliver of the Australian Craton beginning with an arc collision in the Paleocene/Eocene, and then suturing to the main body of New Guinea in the Late Oligocene to Middle Miocene (30-25 Ma, Davies et al., 1997), while the addition of the Accreted Terranes in the northern part of main New Guinea began in the Paleocene in the west (68 Ma) and extended until the Pliocene (5-2 Ma) in the east, the most recent ongoing addition being portions of the New Britain Island arc. The orogeny of the central high mountains is proposed to have occurred in the mid-Miocene (15-11 Ma), together with docking of the Vogelkop Peninsula in the west (Pigram and Davies, 1987; Polhemus and Polhemus, 1998; Quarles van Ufford and Cloos, 2005; Davies, 2012), although Holm et al. (2019) hypothesize that Vogelkop sutured onto the western edge of New Guinea more recently, in the Late Miocene (7-3 Ma). Each of these geological models provides very different spatio-temporal implications for the evolution of biodiversity.

Frogs of the subfamily Asteroprhyinae are a particularly appropriate group to study the role of geologic history on distribution and diversification. Nearly 700 species of microhylid frogs are distributed worldwide across the Americas, Africa, Asia, and Australia (AmphibiaWeb, 2020), but half of them – over 360 recognized species – comprise its largest subfamily, Asterophryinae, which are centered in the Papuan region (New Guinea and its satellite islands, and the Bismarck and Louisiade Archipelagos), and extend into Malaysia, the Philippines, the northeastern coast of Australia, and have recently been described from Thailand and Vietnam. Recent molecular studies indicate they arose during the Miocene, within the past *∼*20 MY (Feng et al., 2017; Hill et al., In Review), coincident with major geological development of New Guinea. High generic diversity (17 recognized genera) and endemism of Asterophryinae relative to the other four native anuran families in New Guinea suggest they were probably the first frog lineage to colonize the region (Van Bocxlaer et al., 2006; Frost et al., 2006; Köhler and Günther, 2008; van der Meijden et al., 2007; Savage, 1973; Rivera et al., 2017; Tu et al., 2018; Hill et al., In Review). Anurans are poor oceanic dispersers, yet a substantial number of species occurring on the offshore islands have deeply divergent sister taxon relationships with lineages on the Papuan mainland (Hill et al., In Review), suggesting either frequent long-distance overwater dispersal or past geologic connection between land masses.

In this study, we reconstruct the biogeographic history of Asteroprhyinae using a densely sampled phylogeny coupled with sophisticated biogeographical and diversification analyses to test alternative geological scenarios. We construct alternative geological hypotheses for (i) slow and steady accumulation of the Accreted and East Papuan Composite terranes, (ii) offshore mobile belt formation, (iii) recent emergence of New Guinea, and (iv) development of the offshore islands which have received little geological study, and allow these hypotheses to compete for the best explanation of range evolution. We infer ancestral ranges and the minimum number of dispersal events to explain the biogeographic distribution of the clade. Furthermore, as alternatives to hypotheses involving plate tectonics, we explore elevational variation and isolation by distance as potential competing hypotheses that might explain the observed patterns of biodiversity.

## Materials and Methods

### The Phylogeny and Geographic Range Data

For our analysis we used the time-calibrated Asterophryinae phylogeny of (Hill et al., In Review), which contains contains 218 tips representing 205 taxa (122 named and an additional 83 putative taxa) across all five tectonic sectors of mainland New Guinea as well as several satellite islands (nine taxa occur at two sites), with two samples each from the Philippines and Sulawesi, Indonesia. Time-calibration was made using widely-agreed upon geological references, specifically the isolation of the islands that form the Louisiade Archipelago (6-4 Ma) and the opening of the Woodlark Basin (6-5 Ma), were used (Hill et al., 1992; Davies et al., 1997; Baldwin et al., 2012; Wallace et al., 2014; Webb et al., 2014), is explained in (Hill et al., In Review; Rivera et al., 2017). Futhermore, the age of the tree is corroborated by Feng et al. (2017), who recovered a 20 M year old age estimate for Asterophryinae using independent data and 20 fossil age constraints from other amphibian groups.

Our samples range over 80 sites across Papua New Guinea and its satellite islands (Fig. 1), spanning a majority of the elevational range of Asterophryne (0-3570 m, Zweifel, 2000, 0-2835m in our dataset). Seventeen sites are located on offshore islands with the remaining 63 occurring on the Papuan mainland. Our geographical sampling includes 13 georegions used to test hypotheses of historical phylogeography: all five recognized major tectonic units of mainland Papua New Guinea: the Eastern Papuan Composite Terrane (E), the Fold Belt (F), the Australian Craton (A), New Britain (B), the Accreted Terrane (A), Misima Island (M), Sudest Island (S), Rossel Island (R), Normanby Island (N), Fergusson Island (G), Woodlark Island (W), the Vogelkop Peninsula (V), and South East Asia (SEA). A summary of the number of species and sites per georegion are listed in Table 1, with metadata for all samples including GPS coordinates being provided in Table 1 of Hill et al. (submitted for publication). We focused our geological exploration on alternative hypotheses of shared geohistory among these regions.

**Table 1.** The number of representative taxa for each georegion included in this analysis. Island area data from (Lyon, 1991) http://islands.unep.ch/Tiarea.htm

| Georegion | Abbrev | Name | # of Taxa | # of Sites | Sq Km |
| --- | --- | --- | --- | --- | --- |
|  |  | **New Guinea** | 159 |  | 785,753 |
|  | **V** | **Vogelkop Peninsula** | 4 | 3 |  |
|  | **A** | **Accreted Terranes** | 32 | 7 |  |
|  | **F** | **Fold Belt** | 19 | 8 |  |
|  | **C** | **Australian Craton** | 3 | 3 |  |
|  | **E** | **EPCT** | 101 | 37 |  |
|  | **N** | **Normanby Island** | 24 | 6 | 1,040 |
|  | **F** | **Fergusson Island** | 3 | 1 | 1,437 |
|  | **B** | **New Britain** | 2 | 2 | 36,520 |
|  | **W** | **Woodlark Island** | 6 | 1 | 874 |
|  | **M** | **Misima Island** | 4 | 1 | 202 |
|  | **S** | **Sudest Island** | 8 | 4 | 866 |
|  | **R** | **Rossel Island** | 8 | 3 | 262 |

**Fig. 1.**
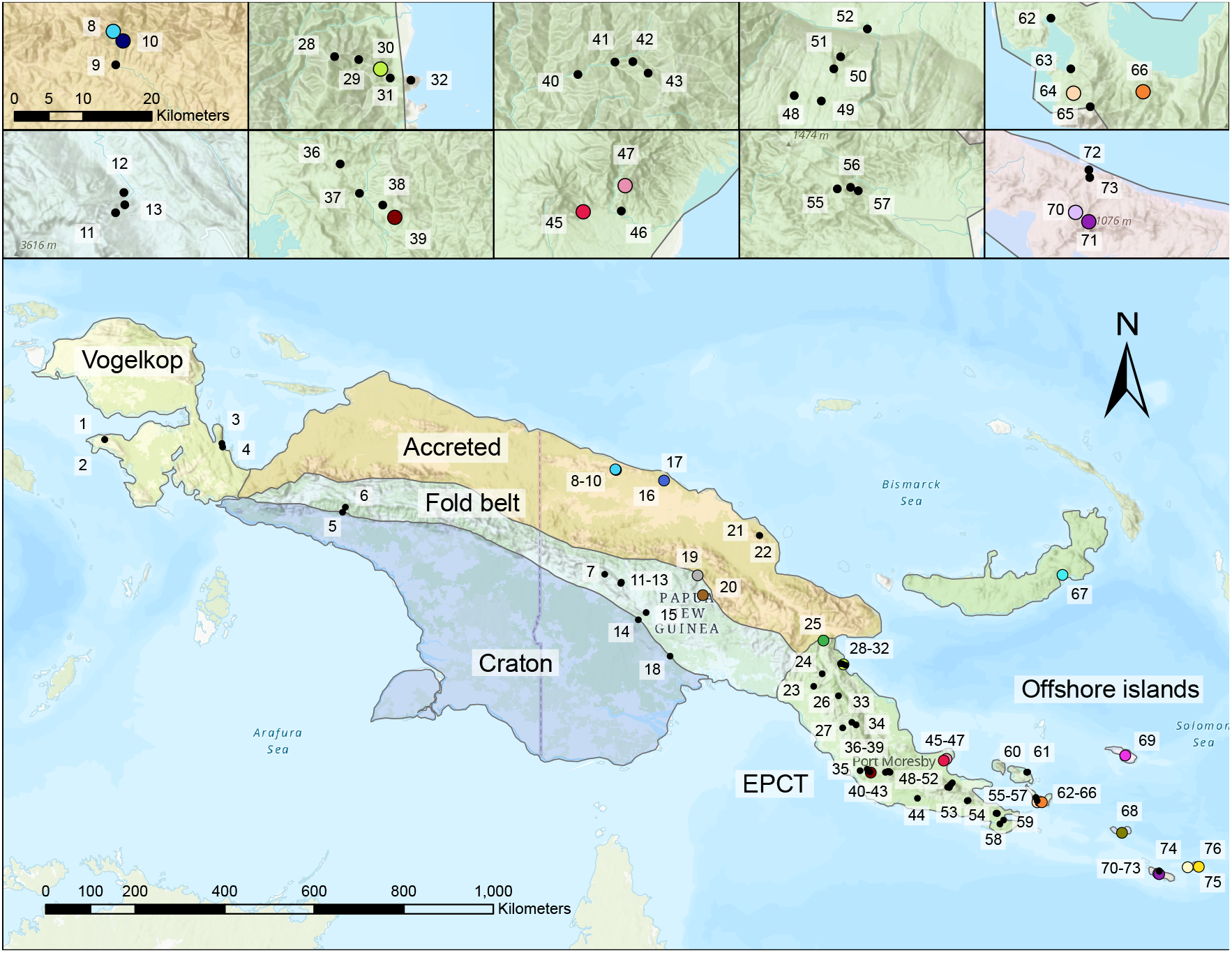
Five major geological regions of NG that illustrate accretion history. The island of New Guinea has a composite history with multiple geologic terranes alamgamating to form the large island. The geologic terranes of New Guinea are labelled following Davies (2012): the Eastern Papuan Composite Terrane, the Accreted Terrane, the Fold Belt, the Australian Craton and the Vogelkop Penninsula. Sampling sites across PNG and satellite islands are indicated by numbers, with insets showing detail for sites that are clustered. Site names, GPS coordinates, and metadata are provided in Table 1 of Hill et al. (submitted for publication).

### Modeling the Evolution of Geographic Range

We modeled the evolution of geographic ranges of Asterophryinae using dispersal-extinction-cladogenesis models (DEC; Ree et al., 2005; Ree and Smith, 2008). The models are described in detail in (Ree et al., 2005; Ree and Smith, 2008), but we briefly review the key components here to describe our application to testing hypotheses of differential dispersal opportunity afforded by alternate geological scenarios. We use “areas” to refer to distinct georegions observed in our dataset, and “range” as the set of areas occupied by a lineage, thus areas are fixed but species ranges can evolve through time. DEC models describe an evolutionary process for biogeographic range evolution along a phylogeny (Ree et al., 2005; Ree and Smith, 2008), and are closely related to stochastic models for discrete character evolution, but importantly differ in breaking up each state change into two possible stochastic events: dispersal to a new area, and extinction from an existing area. Discrete changes from one state to another are assumed to occur randomly through time according to a Markov process with probability matrix **P**_*ij*_(*t*) from ancestor state *i* to descendant state *j*. In matrix form this equation is:

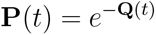

Where **Q** is the instantaneous rate matrix containing both the rates of dispersal *D*_*ij*_ between ancestor range *i* to descendant range *j*, and rates of local extinction *E*_*i*_. For example, for three areas (1, 2, 3, along with the null area *∅* for the possibility of extinction), and with each lineage occupying a maximum of two areas, the possible ranges would be enumerated as *S* = {∅, 1, 2, 3, 12, 13, 23}, and **Q** would be parameterized as:

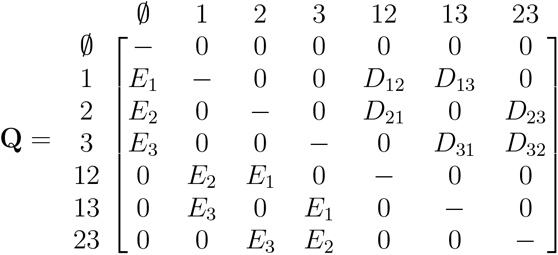

Thus, a frog lineage may expand its range by dispersal into a new area, contract its range when extinction occurs from part of its range, or may change areas if both dispersal to the new and extinction from the old areas occur jointly. These equations model state changes along branches, with nodes assumed to represent speciation events. As the phylogeny is assumed to be true (as with many phylogenetic comparative methods), all possible state combinations at internal nodes compatible with the observed ranges at the tips are integrated over in computing a likelihood for the model. The model outputs include maximum likelihood estimates for the dispersal and extinction rates and inferred ancestral ranges at the nodes. DEC models are flexible enough to allow a variety of evolutionary outcomes, including rapid spread as well as slow biogeographical evolution across a landscape, as one would expect for lineages that are poor dispersers such as frogs, as opposed to free movement from anywhere to anywhere as would be allowed for standard discrete character models.

We focus on testing alternative geological hypotheses that explain the dispersal history of our frogs, implemented by adding constraints that represent hypothetical dispersal barriers. We used the dispersal multipliers matrix in `BioGeoBears` (Matzke, 2014), with “1”s indicating each pair of areas where dispersal is allowed, and “0”s for each pair of areas where dispersal is highly unlikely, as may occur between islands separated by open ocean (we note in practice, the zeros are instead “0.001”s, a small number for the software to return a model fit). The dispersal elements of the Q matrix are multiplied by these constraints, influencing which dispersal paths dominate in the explanation of the data.

We used dispersal multiplier matrices with either pairwise connections or multiway connections to represent our geological scenarios. In particular, New Guinea Island was formed by terranes docking or accreting onto existing terranes along particular margins, which was modeled with a pairwise connection matrix, with the EPCT connected to the Accreted Terranes and the Fold Belt with 1s (brown bars indicating connections in Fig. 2), but not the Australian Craton nor the Vogelkop Peninsula (0s). We note that it is possible for EPCT lineages to reach the Vogelkop, but would require an additional step through the Accreted Terranes, for example, to create a connected path consistent with the difficulty of long-distance dispersal in this system. Alternatively, we have archipelagos which may have been connected at one time, or a hypothesis of islands rifting off from a mainland origin, such as for the Louisiade Archipelago which is hypothesized to have originated as part of the Owen Stanley mountain range of the EPCT prior to its current state as a set of separate oceanic islands. This hypothesis is represented by all members of the affected islands and the EPCT connected by “1”s to each other, forming a block of multi-way connections (green areas of connection in Fig. 2).

**Fig. 2.**
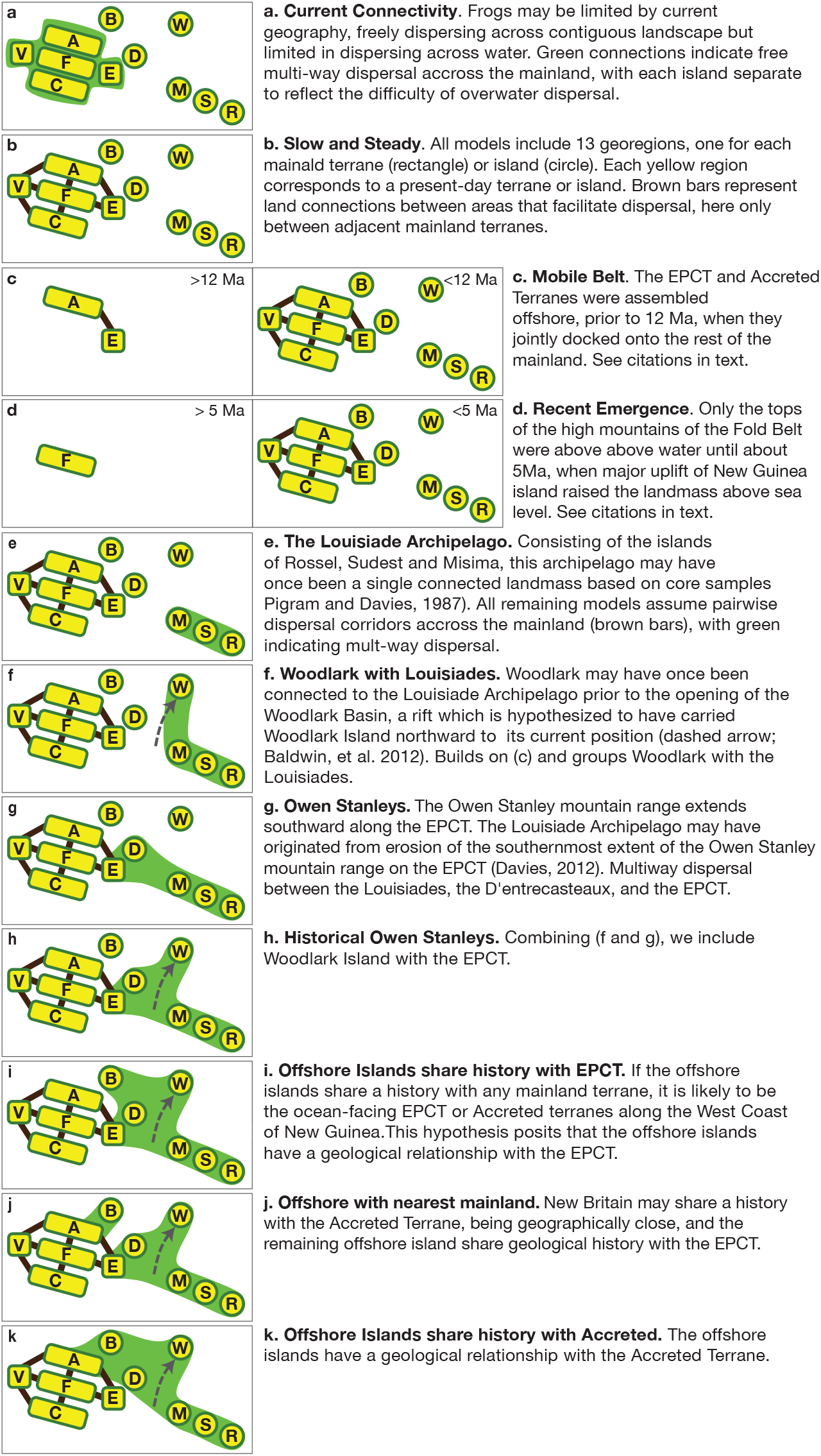
Visual representations of the alternative hypotheses. All models include 13 georegions indicated in yellow, one for each mainland terrane (rectangle) or island (circle). Note that the two islands of the D’entrecasteaux (Normanby and Fergusson) are represented by “D”, and the South East Asian georegion is not shown. Brown bars represent pairwise land connections between areas that facilitate dispersal, whereas areas grouped in green indicate multi-way connections. These alternatives are expressed as modifications to the dispersal multiplier matrix (see methods). Hypotheses **b, c**, and **d** are the major alternatives of slow and steady, mobile belt, and recent emergence, respectively, versus the alternative “null” hypothesis of no geohistory **a**. Hypotheses **e-k** are further refinements of **b** incorporating the history of the offshore islands. Hypotheses **c** and **d** are time-stratified by areas and dispersal corridors available during the periods indicated.

For example, hypothesis **a**, Naive Geohistory, in which only the mainland terrane connections are involved, is represented by the dispersal multiplier matrix below:

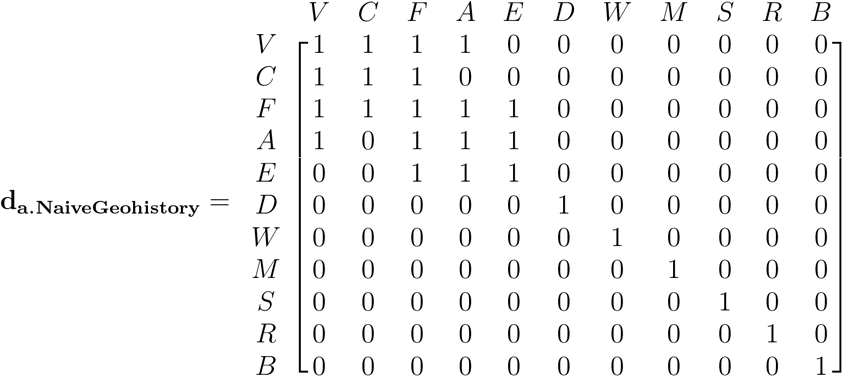

Whereas hypothesis **g** posits that a multi-way connection existed between all offshore islands and the EPCT, but to none of the remaining mainland terranes:

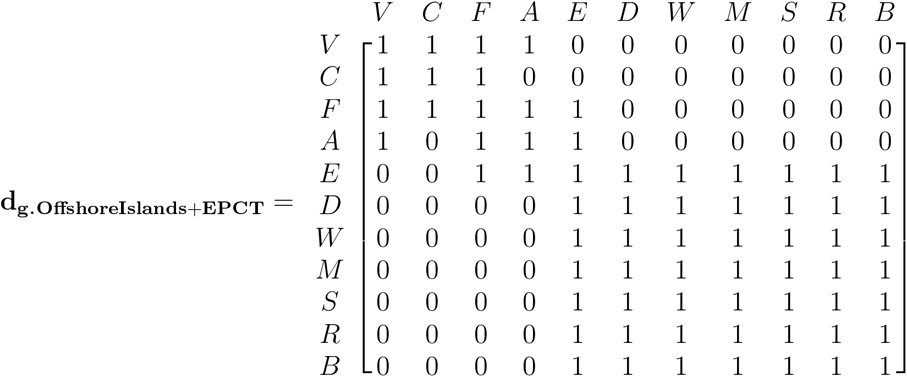

Some of our hypotheses involved recent emergence or assembly of land masses. We used time-stratified models to represent these hypotheses, with the “areas available” matrix and their associated time periods to indicate which areas were available for occupancy during each time strata. A single dispersal multiplier matrix was used across all time strata.

We fit models of historical biogeography using Dispersal-Extinction-Cladeogenesis (DEC; Ree et al., 2005; Ree and Smith, 2008) models as implemented in the R package BioGeoBEARS (Matzke, 2014), allowing a maximum of two areas per taxon. All DEC models have two degrees of freedom for the dispersal and extinction rate parameters. Model comparison was used to evaluate alternative hypotheses of historical biogeography, assessing model fits with the Akaike Information Criterion (AIC; Burnham and Anderson, 2002). Using our best-fitting range evolution models, we estimated ancestral ranges and compared our results with the timings of terrane-accretion and island formation events (Davies et al., 1996, 1997; Pigram and Symond, 1991).

Reconstructed ancestral ranges were assigned to nodes by majority rule, and used to tabulate inferred range shifts, assigning the shift to the ancestral node along which it occurred. Although all Asterophryine species are single-area endemics, we allowed two areas per taxon, as required by the software to return a model fit. We wrote a custom script to apportion the two-area probabilities equally between the original geogregions. For example, if the ancestral reconstruction assigned a 10% probability to a dual geogregion range comprised of the EPCT + Fold Belt, half of the probability would be assigned to the EPCT and half to the Fold Belt. We plotted phylogenies annotated with model results using the R package `ggtree` (Yu et al., 2017).

We note that many biogeographic studies also fit DEC+J models (Matzke, 2014), which allow an additional parameter for “jump” or long-distance dispersal. We found that DEC+J model fits on our data were degenerate as they returned models with zero dispersal and extinction probabilities, apportioning all range evolution to long-distance dispersal. Other authors have reported this model behavior, warning about spurious interpretations especially in situations where all lineages are restricted to single areas, as in our system (Matzke, 2014; Ree and Sanmartín, 2018). We therefore did not use DEC+J models in our study, as these results indicate a pathology of the model fit rather than a reasonable estimate of reality, which would be nonsensical given the poor dispersal abilities of frogs.

### Hypotheses

We tested hypotheses of historical biogeography based on geological history, distance, or elevation as explanatory factors.

#### Geology

Over the past 20M years, there have been significant land movements whose history may leave signatures in the present-day distribution of frog lineages. While it is well established that there are five major tectonic units that joined to form the New Guinea mainland, there are several geological hypotheses regarding the timing and order of their spatial connections. We constructed 11 alternative hypotheses (see Fig. 2 for explanation) to test these ideas. Our null or baseline hypothesis “Current Connectivity”(a) represents opportunities for dispersal based on present-day land connectivity, with no consideration of geological history. Here, mainland frogs can disperse freely across the main island of New Guinea, but overwater dispersal is unlikely. The next three alternative hypotheses represent the major competing ideas for the formation of the mainland: “Slow and Steady”, “Mobile Belt”, and “Recent Emergence” (see introduction for background; and Fig. 2b-d for description).

The remaining hypotheses explore scenarios for the history of the offshore islands and their relation to the mainland. The origins of the Louisiade Archipelago have not been well studied, but based on petrological similarities, they are relatively old, being composed of fore arc metamorphic rocks of at least Miocene age whose protoliths date back to the Cretaceous, and may once have been connected to other land masses, since their metasedimetary rocks are correlative to those in the current Owen Stanley Range of the Papuan Peninsula (Fig. 2e, Davies and Smith, 1971; Pigram and Davies, 1987; Baldwin et al., 2012). Prior to the opening of the Woodlark Basin, Woodlark Island was in close proximity to the Louisiade Archipelago (Fig. 2f, Pigram and Davies, 1987), possibly representing an element of the volcanic back-arc behind the Louisiade fore-arc (Webb et al., 2014). In its present-day position, Woodlark has been significantly displaced northward from its paleo-position, due to the rapid opening of the Woodlark Basin around 5 Ma. Another correlated hypothesis regarding the mountains of the Louisiade Archipelago is that they may represent the southern most extent of the ancestral Owen Stanley mountain range, which might once have been contiguous within the EPCT before its southeastern extension was isolated due to erosion and subsidence (Fig. 2g,h). This idea is again supported by petrological similarities (Davies and Smith, 1971; Pigram and Davies, 1987). We also include the D’Entrecasteaux Islands because of their very close proximity to the EPCT (although three islands comprise the D’Entrecasteaux group – Normanby, Fergusson, and Goodenough, we have frog data from only the first two, so our subsequent discussions treat this unit as a two-island complex). The final hypotheses test whether the offshore islands share a history with either the EPCT or the Accreted Terranes, or both, as the latter lie on an active margin in proximity to the offshore islands (Fig. 2i-k).

#### Island Distance

The islands of the Bismark, D’Entrecasteaux, and Louisiade groups lying off the northeast coast of New Guinea are surrounded by waters over 100 m deep and vary in distance to the mainland, which in turn should be inversely proportional to the probability of overwater dispersal. Therefore, absent any influence of geological history, nearby islands, such as New Britain and the D’Entrecasteaux islands (*<*100km away from the mainland), are expected to experience greater rates of dispersal than distant islands such as the Louisiade Archipelago (*>*200km). We tested this idea by weighting the dispersal multiplier matrix by distance categories. Thus, the nearby D’entrecasteaux Islands (Normanby and Fergusson) and New Britain in relation to any mainland region are assigned a dispersal multiplier of .1, whereas Woodlark Island and the islands of the Louisiade Archipelago are assigned a dispersal multiplier of .001. Within-archipelago movement is assumed to be unconstrained (i.e., within the D’Entrecasteaux islands, within the Louisiades + Woodlark), with a dispersal multiplier of 1.

#### Elevation

We also tested for evidence of elevational specialization by binning the ranges used in DEC models according to elevation. One possibility is that frogs are adapted to a particular elevational range and may only disperse to new regions within the same elevation. Alternatively, we may see patterns of a gradual or stepwise shift in elevational ranges as the frogs slowly ascend or descend a mountain. We selected elevation categories empirically based on the distribution of observed elevation ranges, with limits chosen to create bins containing roughly equal proportions of frog species: 0-200m, 200-550m, 550-900m, 900m-2000m, 2000m+. We applied both the unconstrained DEC model and a DEC model with dispersal constraints connecting only adjacent elevational ranges. Because these are different data than the georegion data, we cannot directly compare the fit of elevation models to geological or distance models, however, we can compare across elevation models. Violin plots of the elevational distribution were created using the `ggplot2` package in (R Core Team, 2020).

## Results

Because of the multiple hypotheses regarding the mainland of New Guinea and the offshore islands and their potential combinations, we took a two-tiered approach to evaluating our many moving parts, first evaluating the major alternatives regarding the mainland, followed by the refinements involving the offshore islands. Among our mainland-centered hypotheses, the distribution of Asterophryinae is better explained by the “b: Slow and Steady” hypothesis which was far superior to the “c: Mobile Belt” and “a: Current Connectivity” hypotheses by 53 and 64 AIC units, respectively. “d: Recent Emergence” provided a subtantially worse fit, by 308 AIC units. However, “Slow and Steady” was vastly improved by incorporating the geological history of the offshore islands.

The best-fitting overall hypothesis was “j: Offshore with Nearest Mainland”; which built upon “Slow and Steady” by facilitating dispersal to the offshore islands from their nearest mainland units (the Accreted Terranes for New Britain, and all others connected to the EPCT; Table 2), and was far superior to all other models. The second best model, by 6 AIC units, was hypothesis “i” in which all of the offshore islands including New Britain are connected to the EPCT (and not the Accreted Terranes). Switching the shared history of the offshore islands from with EPCT to with the Accreted Terranes provided a very poor fit, with a ΔAIC of 188 for hypothesis “k”. Overwater distance provided a poor explanation of the data with a ΔAIC of 118. Taken together, these results indicate that with the exception of New Britain, all of the offshore islands share a history with the EPCT.

**Table 2.**
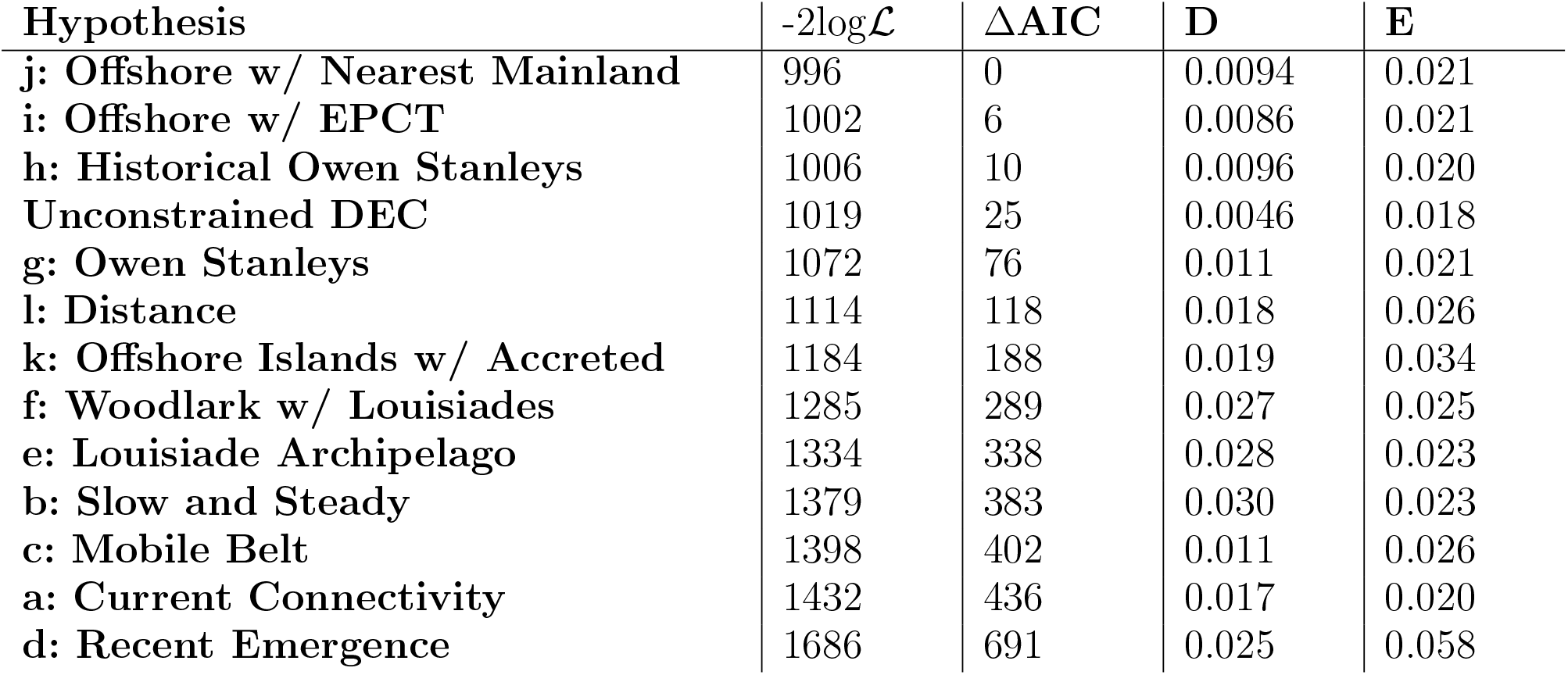
Performance of alternative hypotheses for geographic range evolution. For each hypothesis, the likelihood values (-2log*ℒ*) and difference in Akaike Information Criterion from the best fitting model (ΔAIC) are given along with parameter estimates for the rate of dispersal (D) and extinction (E). Degrees of freedom for all models is 2.

Asterophrine frogs occur across a wide elevational range (0-3570 m, Zweifel, 2000, 0-2835m in our dataset), concentrated in the mid-low elevation ranges (mean 813m, median 600m, Fig. 3A). Many well-represented genera have relatively even distribution across a wide elevation range, which mirrors the subfamily-wide distribution (Fig. 3B). Accounting for elevation in DEC models did not reveal any consistent pattern with regard to diversification. Constraining elevational shifts to adjacent categories provided a much poorer fit by 81 AIC units than the unconstrained elevation model, indicating frequent shifts between non-adjacent elevational ranges, which is evident in the reconstruction of the ancestral states (see Supplementary Fig. S1). Indeed, multiple elevational ranges can be found in any clade with very little phylogenetic conservation of elevational ranges.

**Fig. 3.**
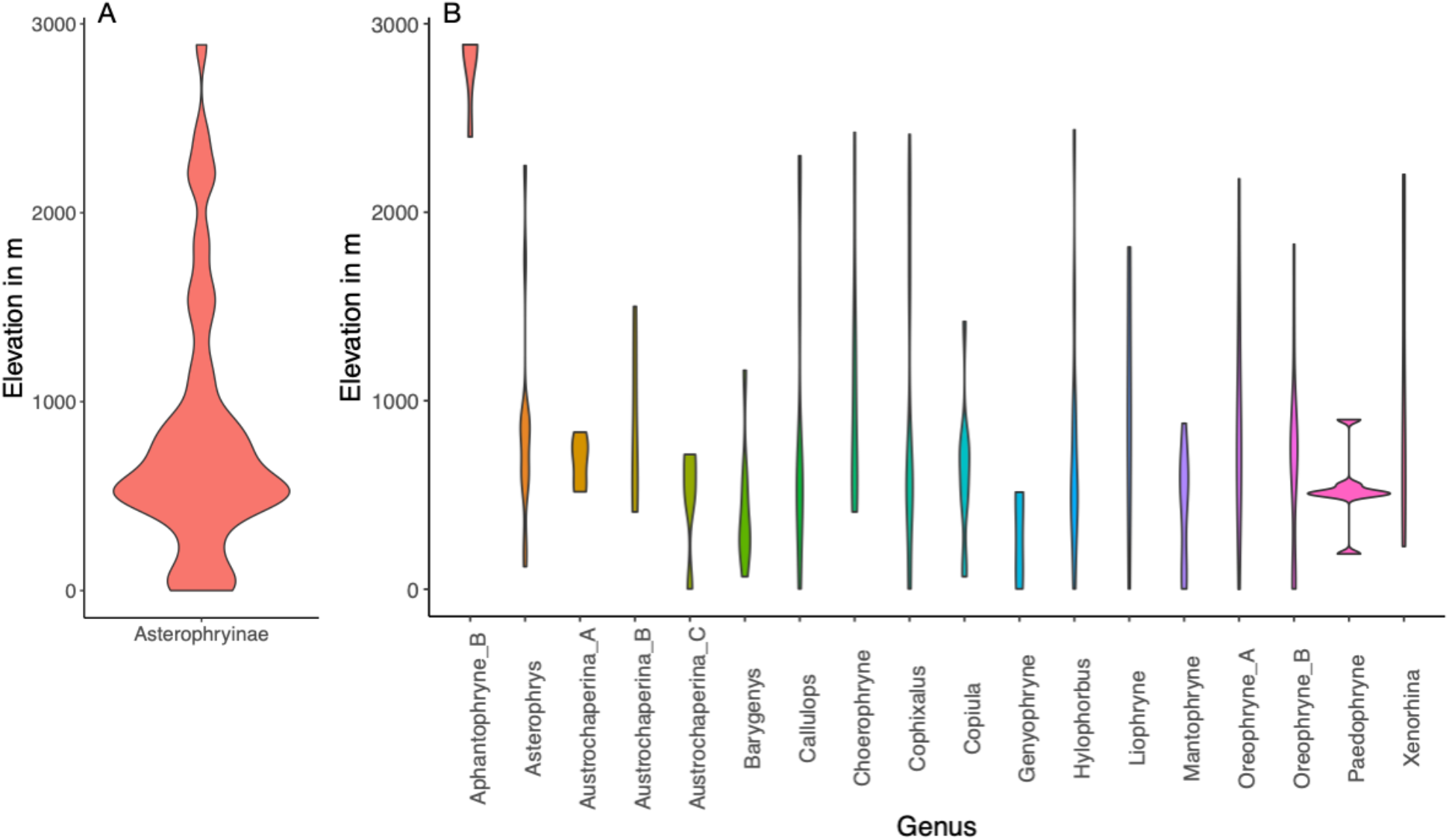
Distribution of Asterophrinae across elevations across the subfamily (A), and separated by genus (B). Width of violin plots is proportional to the density of species across elevations.

### Patterns of Dispersal

From the best-fit model we are able to reconstruct the history of Asterophryinae dispersal and evolution across New Guinea (Fig. 4). The MRCA to Asterophryinae most likely arrived on the EPCT, approximately 20 Ma. Most genera originated in the EPCT, and in general, clades show limited dispersal, tending to remain within a georegion (Fig. 4). However, there are several genera that have had parallel dispersal followed by diversification events from the EPCT to the Accreted Terranes, or from the EPCT to the Fold Belt (Fig. 5), with a burst of these diversifying dispersals approximately 17-15 Ma. The clade diversified and dispersed across the region in 71 dispersal events condensed into four major periods: 20-15 Ma, 15-10 Ma, 10-3 Ma and *<* 3 Ma. The dispersals to the islands in nearly all cases occurred from the EPCT, the one exception is a single dispersal from the Accreted Terrane to New Britain. Generally, as time moved toward the present, the numbers of dispersals accelerated until about 3 Ma (Table 3). Between 3 Ma and the present, dispersals steeply declined, with the majority of dispersals between the Papuan mainland and the offshore islands, and with only two dispersals occurring between tectonic units within the Papuan mainland.

**Table 3.**
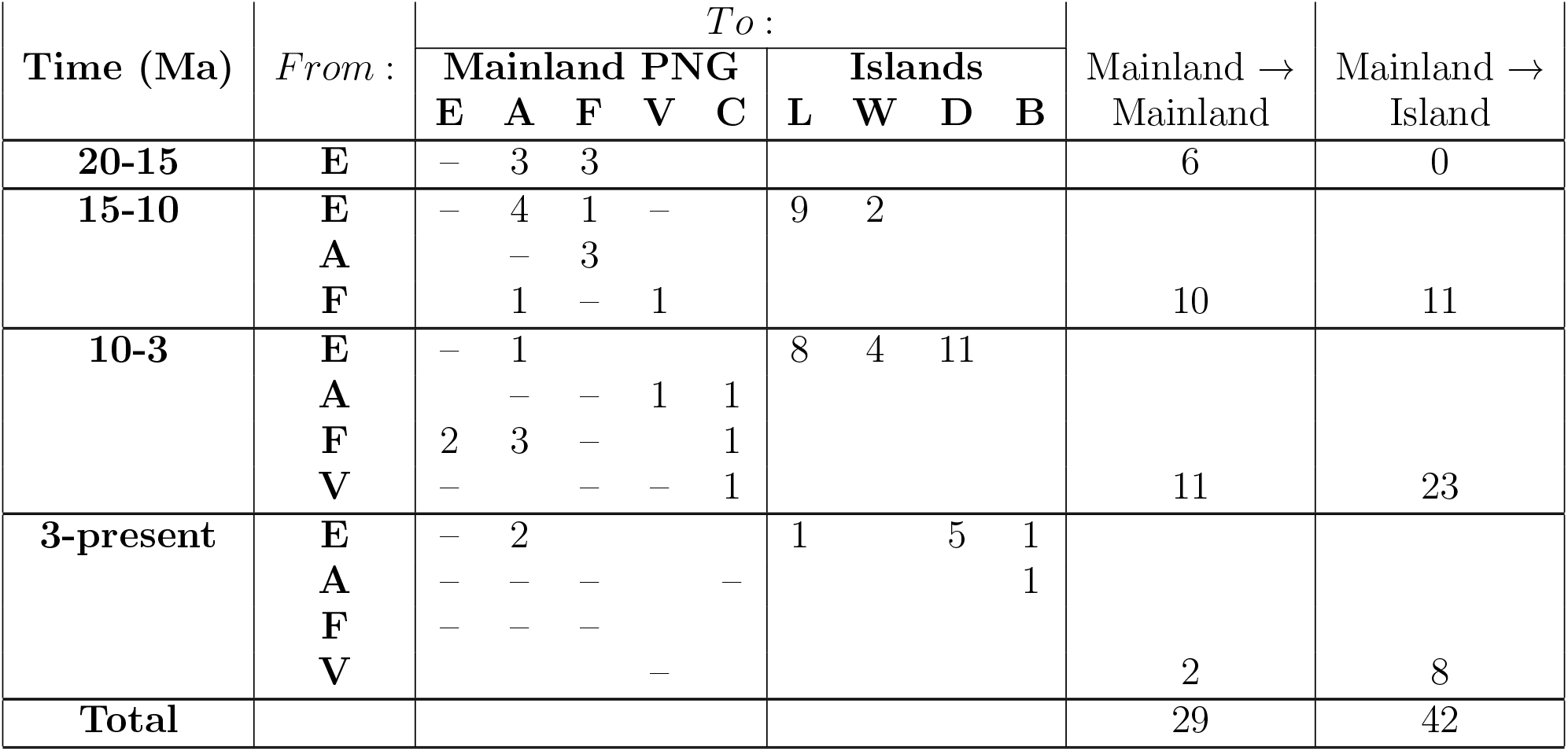
Numbers of Asterophryinae dispersals through time between mainland and offshore islands of New Guinea. Mainland Terranes: E = EPCT, A = Accreted Terrane, F = Fold Belt, V = Vogelkop Peninsula, C = Australian Craton. Islands: L = Louisiade Archipelago, W = Woodlark Island, D = D’entrecasteaux Islands, B = New Britain Island.

**Fig. 4.**
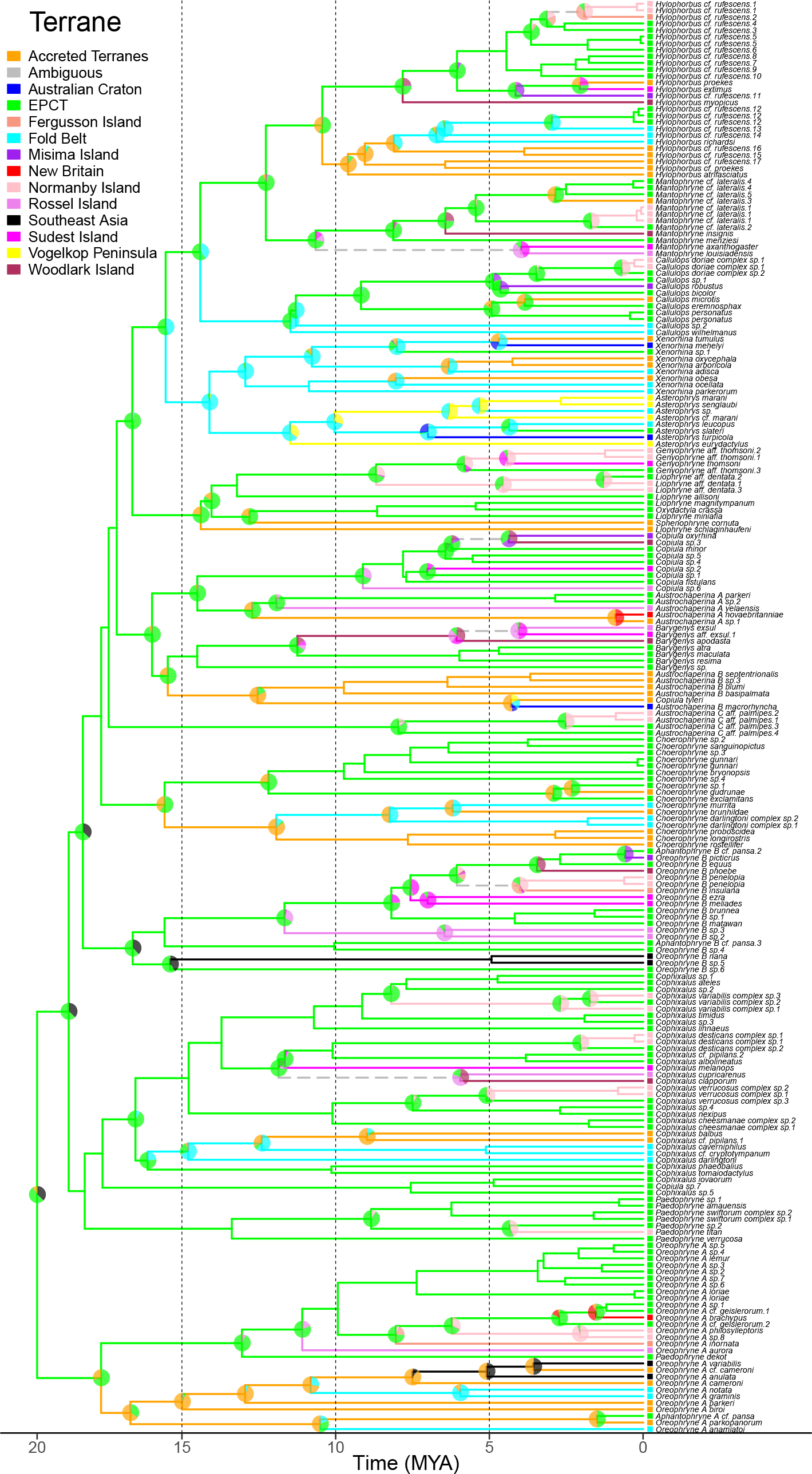
Reconstruction of Asterophryinae dispersal across New Guinea under the best fit DEC model. As all species are single-georegion endemics, probabilities are apportioned to single georegions.

**Fig. 5.**
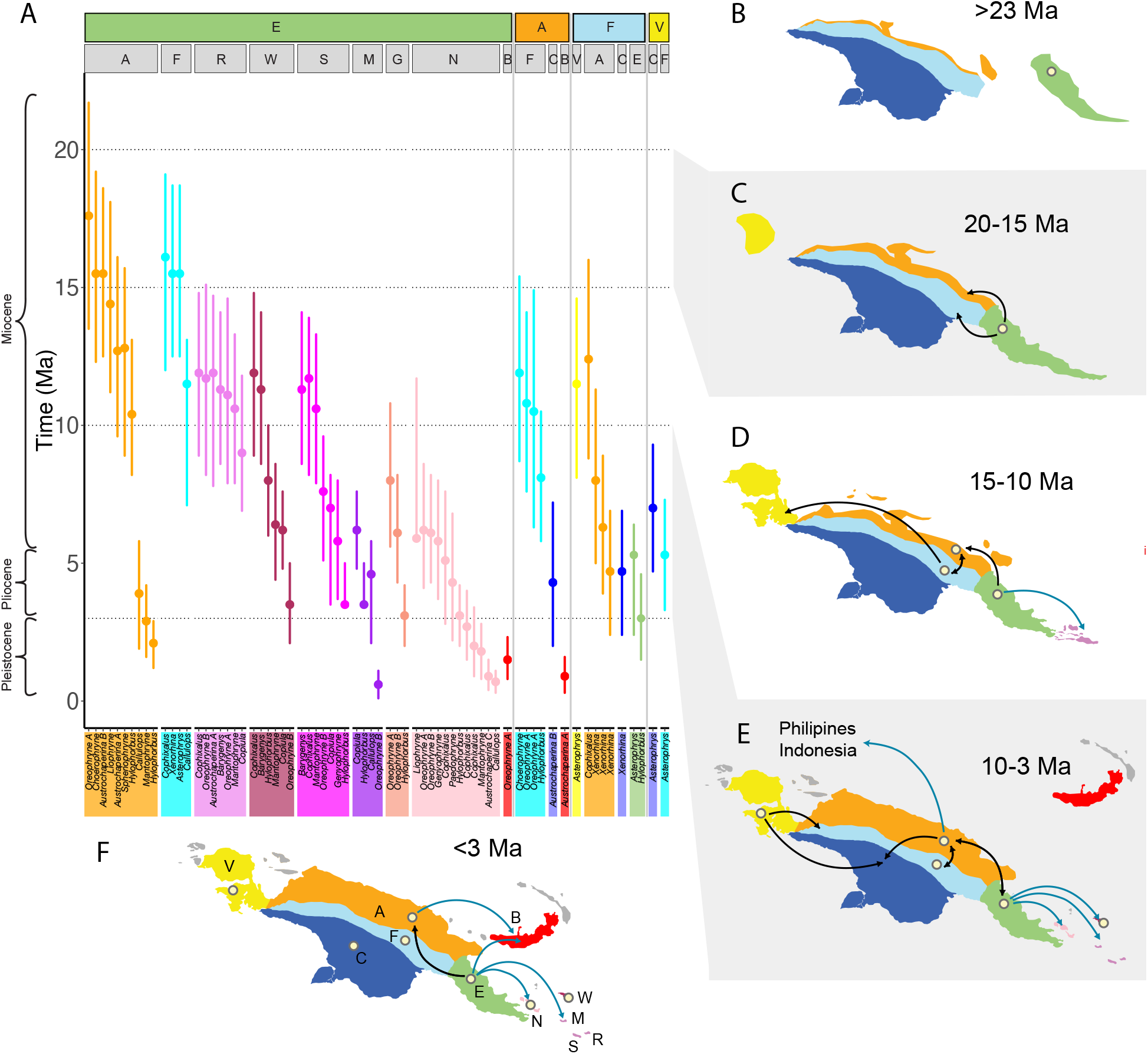
The dispersal of Asterophryine frogs across New Guinea between terranes through time inferred from our phylogenetic analysis and the corresponding geological evolution. (A) Timings of independent dispersal events between terranes or islands inferred from DEC analysis (Fig 4). Genera involved are along the lower X-axis. The source terrane is indicated along the first row of the upper X-axis, arrival terrane or island along the second row. Points and genera blocks are colored by the arrival terrane/island. (E=EPCT in green, A=Accreted terranes in orange, F=Fold belt in light blue, and V=Vogelkop peninsula in yellow, R=Rossel Island in lilac, W=Woodlark Island in maroon, S=Sudest Island in fuscia, M=Misima Island in eggplant, G=Ferguson Island in coral, N=Normanby Island in rose, B=New Britain Island in red). (B) - (F) Corresponding land movements and the evolving islands. Beige dots indicate terranes/islands with frogs established at the start of the epoch. (B) Prior to 23 Ma, the EPCT is approaching the nascent New Guinea island bringing the ancestors of Asterophryinae. (C) 23-20 Ma, the EPCT has docked onto the growing mainland, facilitating overland dispersal to the Accreted Terranes and the Fold Belt. (D) 15-10 Ma, the Vogelkop Peninsula docks, allowing dispersal. Dispersal continues between mainland terranes, on the southern tip of the EPCT the first genera disperse to Rossel, Woodlark, and Sudest islands as they break away from the Owen Stanley mountain range, encouraging speciation. (E) 10-3 Ma, the Accreted Terranes have grown along the north coast, ongoing dispersal and speciation across land bridges and short water gaps until islands move further away. (F) The D’Entrecasteaux islands (Normanby, Ferguson included in this study) have emerged very close to the mainland, and New Britain is on a collision course toward New Guinea, with the first dispersals to these islands.

## Discussion

The small, terrestrial Asterophryine frogs have a surprisingly wide distribution across the Papuan region. We recover the EPCT as the center of origin of Asterophryinae placed at about 20 Ma (Feng et al., 2017; Hill et al., In Review), with subsequent spread throughout mainland New Guinea and across satellite islands, some more than 200km away. Furthermore, phylogenetic analysis of range evolution demonstrates that they have criss-crossed across large distances over land and open ocean at multiple times throughout their history. From our DEC model, we recover an overall rate of extinction that is higher than the rate of dispersal, consistent with a clade of low-dispersing organisms that tend to readily fragment by isolation. Our analyses do reveal some macroevolutionary patterns of isolation by distance, with some sub-generic clades found predominantly within a single terrane, but what is unexpected is the frequent pattern of apparent dispersal across the phylogeny, requiring a minimum of 71 events that involve some sort of long-distance traversal (Table 3). If we consider that a tree with 218 taxa has 436 branches which could possibly recover a range transition event, the Asterophryinae phylogeny reveals a major geographical shift over 16% of its branches. In this paper we illustrate the power of the hypothesis-comparison approach for explaining complex distributional patterns, allowing autecological and geological ideas to compete for the best explanation of the data. We considered three possible explanations for this overall distributional pattern: (1) specialization to elevation as is common for many New Guinea taxa, (2) long distance dispersal and subsequent diversification, and (3) conveyance via plate tectonics, of which we have several sub-hypotheses involving both the timings and spatial dynamics of the mainland tectonic units and the offshore islands. As we now show, the first two provide little explanatory power, whereas the third explains most of the data and requires only rare long-distance dispersal.

### Elevation

We do not find any signatures of elevational specialization across the subfamily Asterophryinae. The most common elevation range early in the history of Asteroprhyinae *∼*20-15 Ma is the low-mid elevation range (200-550m, Fig. 3, Supplemental Fig. S1). From there, they have frequently shifted elevation ranges, sometimes with large jumps in elevation. Current species diversity is not concentrated at high elevation, either as a whole or by genus (Fig. 3; in contrast with Oliver et al., 2017), nor do they show any tendency to move into or out of high elevation habitats. While the earliest shift to high elevation appears roughly coincident with orogeny of the central cordillera and development of altitudinal variation in New Guinea at *∼*11Ma, upward shifts subsequently occur sporadically from the time of orogeny to the present throughout the subfamily. We note that our data shows *Asterophrys* as a high elevation genus, but as the monophyly of this group is in question, we remain skeptical about this result (Hill et al., In Review). Oliver et al. (2017) reported montane endemism in a subclade of the genus *Choerophryne* along the central cordillera, with a second mid-elevation clade along the north coast (our Fold Belt and Accreted Terranes, respectively). We do not find conflict with this pattern, but our dataset includes a large clade of low-mid elevation species from the EPCT that are not well-represented in the Oliver et al. (2017) study, and thus we do not find unusual montane occupancy. Rather, what we find more striking is the large number of genera (including *Choerophryne*) that have extremely broad elevational ranges. What was probably not appreciated prior to Hill et al. (In Review) is that Asterophryinae possess a high degree of phylogenetic structure at surprisingly small spatial scales. Therefore, as elevation is inextricably linked to geography, montane endemism in a subclade with low dispersal could simply result from geological evolution. Furthermore, high altitude shifts, where they do occur, tend to lead to terminal taxa so that none of the shifts give rise to in-situ diversification (Supplemental Fig. S1), as one might expect if changes in elevation open ecological opportunity. Thus, we do not find any evidence that elevation is a driver of evolution in this clade.

### Dispersal across the ocean

We recovered no fewer than 42 connections between the mainland and the offshore islands, which can contain complex communities of Asterophryinae despite being small islands more than 200 km away. All but one of our offshore island species have sister-taxon connections to the EPCT, and similarly speciose assemblages irrespective of island distance (e.g, the close D’entrecasteaux islands and the far Louisades). If dispersal was trans-oceanic, it would imply that more than half of the movements were across open ocean. Historically, amphibians have been regarded as excellent indicators of vicariance (Duellman and Trueb, 1994), as adult anurans suffer toxicity at 80% of ocean salinity, and embryos are developmentally impaired at only 25% of ocean salinity (Gordon et al., 1961; Hopkins et al., 2012; Seymour, 1994). One potential explanation is that frogs may raft on fallen trees, partially partial protected from the ocean, yet to reach the offshore islands from the mainland of New Guinea, frogs must travel south-to-east. However, dispersing frogs would be buffeted by ocean currents in the opposite direction, and for five months of the year, these are strengthened by southeast monsoon winds (McGregor et al., 2008; McAlpine et al., 1983). Nevertheless, we found that Asterophryine frogs reached at least three islands that have never been connected to any landmass: Fergusson and Goodenough islands of the D’entrecasteux archipelago, which are small, nearby, and recently uplifted islands, and contain three and seven species of Asterophryne. New Britain is a rather large island, but contains only two species (Table 1). These examples provide clear evidence that overwater dispersal can occur, although it is rare (de Queiroz, 2005; Vences et al., 2003).

If we assume that the islands have always been separated by fixed distances, as would be the case for hotspot island archipelagos such as the Hawaiian islands, the predictions of island biogeography theory do not hold. New Britain, the largest satellite island has the fewest species despite being relatively close, and tiny islands contain many species irrespective of distance. We found no support for island distance providing a general explanation of the data (distance model ΔAIC = 188,Table 2). However, we can begin to see a role for dispersal if geological history is taken into account. Frogs have only recently dispersed to New Britain, which is situated on the Bismarck plate and has been traveling south and west toward New Guinea and is therefore presently as close to New Guinea as it has ever been, which may be at the maximum range for oceanic dispersal. For the remaining islands, the overall pattern can make sense if dispersal was not trans-oceanic (see below).

### Movement of the Land

Our results show the tremendous influence that geological history has had in the distribution and generation of biodiversity in Asterophryinae. We found support for five mainland and eight offshore georegions that over the past 20 MY, have joined together, rifted apart, and moved across the surface of the earth to provide ample opportunity for dispersal to new lands and evolution by vicariance (see summary in Supplementary Fig. S2). This dynamic landscape provided opportunity for at least 66 transitions between mainland terranes, or between the mainland and offshore islands excluding 5 cases of oceanic dispersal in our dataset to New Britain and Fergusson (see below for more detail). We recovered the EPCT as the center of origin of Asterophryinae, with the ancestor to the clade arriving on this terrane approximately 20 Ma. The EPCT has also acted as the primary source for Asterophryinae dispersal. The timings of diversification and ancestral ranges correspond well to some models of geological history that describe the formation of New Guinea as a slow but steady stream of accretion events. Furthermore, viewing the dispersal of Asterophryinae through a geological lens, we see that Asterophryinae did not have to repeatedly and independently disperse across and large ocean barriers to the offshore islands in the majority of instances, but that currently observed distributions may have been predominantly generated through vicariance and short-distance oceanic dispersal as historical land connections disappeared and islands slowly became separated from each other.

### The difficulty of long-distance dispersal over land

A model limiting dispersal to adjacent tectonic units fared substantially better than an unrestricted one allowing free dispersal between any two mainland units (“Slow and Steady” fit better than “Current Connectivity” by 53 AIC units). This can be interpreted as the difficulty of long-distance jumps, for example, we do not see any instances in our data of frogs directly dispersing from the EPCT to the Vogelkop Peninsula. Nearly all range shifts occur in a single step (Fig. 5). Where a lineage passes through multiple terranes, they do so in a stepwise manner: the ancestor of *Asterophrys* and *Xenorhina* originated on the EPCT, then dispersed to the Fold Belt, then to the Vogelkop peninsula. We observe only one multiple terrane traversal, from the Accreted Terranes to the Australian Craton. The model indicates that long-distance overland dispersal is possible, but is also rare.

### The EPCT is the center of origin and ongoing diversification

The EPCT is a known center of endemism and species richness. Based on cicada relationships, Duffels (1983) concluded that the Papuan Peninsula (EPCT) was a separate biogeographic area, and not merely an eastward extension of the New Guinea central mountains, and (Boer and Duffels, 1996) identified it as a discrete area of endemism within the Melanesian region. Kalkman et al. (2018) showed that the EPCT harbored a large number of endemic damselflies and concluded that a range expansion out of the EPCT gave rise to the rest of a species-rich clade of damselflies now found in New Guinea. This is plausible if they arrived from the EPCT to find much ecological opportunity on a still relatively depauperate and more recently uplifted New Guinea. Similarly, Polhemus and Polhemus (2004) noted that the EPCT support an unusually rich biota of veliid water bugs representing numerous endemic genera and species not found in the remainder of New Guinea as a whole, while at the same time lacking certain endemic genera occurring in the main body of the island indicating separate histories of faunal evolution followed by fusion of the two land masses. Kraus (2021) proposed that the opening of the Woodlark rift and its ongoing extension has created a series of vicariance events promoting speciation in multiple lineages. A further possibility is that as a composite terrane, the accretional activity may have brought communities together via collision, or at least within proximity so that dispersal to islands is possible.

The concept of a composite land mass in eastern New Guinea formed by the collision of an Australian continental fragment and an island arc was advanced over 40 years ago (Pieters, 1978; Hamilton, 1979), and has been subsequently supported by more recent research (Webb et al., 2014). Specifically, the basement rocks of southeastern New Guinea (the nascent EPCT) represent a displaced fragment of the Australian continental margin rifted away during the Cretaceous and moved northeastward by subsequent Paleocene-Eocene seafloor spreading (Zirakparvar et al., 2013). This fragment subsequently collided with a Late Paleocene-Early Eocene island arc to form the initial core of what would become the Papuan Peninsula (Davies and Warren, 1988), leading to the emplacement of a broad ophiolite belt banked against the northern margin of this accreted unit. Such a scenario would imply that emergent land masses linked to island arcs may have been present in the vicinity of the current EPCT as early as the Paleocene, and that orogeny in the EPCT preceded that in central New Guinea by at least 10 My (Quarles van Ufford and Cloos, 2005), making it one of the earliest high emergent land masses in the region.

Our studies support this geological scenario, indicating that the EPCT is not only the origin of diversity for Asterophryinae, but a source for ongoing diversification and outward expansion. Asterophryinae originated on the EPCT at least 20 Ma, as did 15 of the 18 genera (this study, Hill et al., In Review; Rivera et al., 2017; Feng et al., 2017), and began to disperse to the Fold Belt or the Accreted Terranes beginning around 17 Ma (Fig. 4, 5). We see that the vast majority of dispersals come out of the EPCT. Of the mainland to mainland dispersals, 48% originate from the EPCT, and dispersals to islands are nearly exclusively from the EPCT at 98%. This strong signal for an origin on the EPCT refutes the “Mobile Belt” hypothesis, which would predict a blend of EPCT and Accreted Terranes (*∼*20-17Ma), and instead lends strong support to the “Slow and Steady” geological model of Pigram and Davies (1987); Davies (2012), a key prediction of which is that the Papuan mainland, except for the northern coast of the Accreted Terranes, was subaerial prior to the early Miocene, with the EPCT docking onto the growing New Guinea composite land mass during the Late Oligocene to Middle Miocene (Davies et al., 1997). The dominance of the clade at low-mid elevation in the early Miocene, as well as the exceptionally poor fit of the “Recent Emergence” model rejects the hypothesis of Hall (2002) that the bulk of the New Guinea mainland was submerged until 5 Ma.

What is more remarkable, however, is the coincidence of the timings between phylogenetically independent dispersals across multiple genera and a number of hypothesized geological events. First, the docking of the EPCT to the Paupuan mainland would have created new overland dispersal routes between major landmasses from the EPCT to the Accreted Terranes and the Fold Belt, bringing the first frogs to the growing New Guinea island. Indeed in our biogeographic results, we see a wave of dispersal across the older genera of Asterophryinae in an “out of the EPCT to the north” pattern in: Oreophryne A, *Cophixalus, Choerophryne*, and *Austrochaperina B* in relation to *Baryengys* (Fig. 5), with each genus split into EPCT vs. Accreted Terrane/Fold Belt divisions at the same point in Asterophryine history, leading to major subclades with ongoing diversification.

About 12 Ma, there are multiple major geological events: orogeny of the central cordillera, docking of the Vogelkop, and the opening of the Woodlark basin. There is a consensus amongst many geologists that the orogeny of the central mountains of the Fold Belt began about 12 Ma (Pigram and Davies, 1987; Pigram and Symond, 1991; Davies et al., 1996; Quarles van Ufford and Cloos, 2005; Davies, 2012; Baldwin et al., 2012). Asterophryinae dispersal from the EPCT into the Accreted Terranes and/or the Fold Belt started around 17 Ma and ending *∼*12 Ma (Fig. 5), when the central mountains may have risen high enough to become a geographic barrier for some lineages. A subset of genera dispersed from the Accreted Terranes to the Fold Belt *∼*12-8 Ma, as the mountains were rising. A final burst of dispersal of three genera from the EPCT to the Accreted Terranes occurred *∼*4 Ma, with the docking of the Huon-Finisterre-Adelbert blocks of the Bismarck Plate, which must have arrived without any preexisting Asterophryinae species.

Contemporaneously in time, Asterophryinae expansion to the Vogelkop Peninsula began *∼*14 Ma from the Fold Belt, which is also coincident with the timing of the docking of the Vogelkop Peninsula (Pigram and Davies, 1987; Polhemus and Polhemus, 1998; Quarles van Ufford and Cloos, 2005; Davies, 2012), rather than the later date of 7-3 Ma proposed by Holm et al. (2019). The timing of these dispersals closer to the mid-Miocene further supports the “Slow and Steady” model rather than the “Recent Emergence” model (Hall, 1998, 2002).

Along the southern tip of New Guinea, also beginning about *∼*12 Ma, was the rifting off of the southern end of the Owen Stanley Mountain range in the EPCT into the Louisiade archipelago and Woodlark Island. We see a very strong signature of this event in our model fits, as any model linking the Louisiades and Woodlark to the EPCT provides a far superior fit to similar models without this motif. At this time, the EPCT would have presumably been fully populated with potential source populations for dispersal across small water gaps or may have already populated the area that became newly isolated islands. This scenario would readily explain the skewed distribution of dispersals from numerous genera coincident in time to these islands, concentrated at the 12 Ma mark, and tapering off as the islands rift farther away. It would also explain why the communities on these islands were seeded from the EPCT, and not arising by rare dispersal followed by in-situ diversification, and also explain why species do not disperse in a stepping stone manner to the islands. Indeed it is remarkable that all species have closest sisters on the EPCT, and in-situ radiation on the islands is rare.

Finally, the recent rise of the D’Entrecasteaux islands demonstrate that at very close distances, oceanic dispersal is possible even for terrestrial frogs, as lineages on these nearby islands which were never connected are related to sisters on the EPCT. The best-fitting overall hypothesis was “j:Offshore Islands with Nearest Mainland”, which is a refinement of “Slow and Steady” and supports these geological scenarios regarding the origins of the offshore islands discussed below. Thus, the hypderdiversity of this low-dispersal clade can be explained by major land movements that provide multiple vicariance events through time, which allowed numerous genera to simultaneously speciate (Figure 5), and lend strong support for the role of geology in driving ongoing diversification.

### The offshore islands

Originally, we thought that our phylogeny (presented in: Hill et al., In Review) was wrong. How could small terrestrial frogs disperse over hundreds of kilometers of open ocean, separated by 10 MY from their nearest sister lineage on the EPCT? We did not find that frogs dispersed in a stepping-stone fashion from nearer to more distant islands. All source populations are directly from the mainland. The answer that emerges from the amalgamation of evidence is that the frogs did not swim, but dispersed over land connections or very narrow water gaps, which themselves evolved over time.

### The Louisade Archipelago

These islands are an excellent case in point, being home to over 18 candidate species of Asterophryinae (Boulenger, 1890; Mé helÿ, 1901; Zweifel, 1963, 1972; Zweifel and Tyler, 1982; Burton, 1986; Zweifel, 2000; Richards and Oliver, 2007; Kraus and Allison, 2009; Kraus, 2016), yet separated by over 200km of saltwater barrier from any potential source population. Rather than inferring 18 independent dispersals within the last 12 million years, an alternative hypothesis is that they may have fragmented from the Papuan mainland. If dispersal was to a remote oceanic island at fixed distance from the source population, we would expect to see a random pattern in the timing of dispersal events. Instead, the data show that the dispersals, all originating from the EPCT, are highly concentrated at *∼*12 Ma, followed by a tapering off until *∼*4 Ma, consistent with islands fragmenting off of a mainland source (Fig. 5). Geological evidence supports this scenario. Petrological samples taken from the Louisiade Archipelago are similar in composition to those of the Owen Stanley Range on the EPCT indicating a shared geological history between these two seemingly disjunct landmasses (Pigram and Davies, 1987; Davies, 2012; Baldwin et al., 2012).

In the work of Pigram and Davies (1987) the Louisiade Islands are shown as a disjunct sector of the Owen Stanley Terrane, most of which lies in the interior of the Papuan Peninsula. In their current form, they represent the partially drowned remnants of an old mountain range that ran down the middle of what is now the EPCT, and the metasedimentary rocks of the Louisiades have been correlated with those of the Owen Stanley Range in the Papuan Peninsula (Davies and Smith, 1971). The protoliths of these metasedimentary rocks are Cretaceous volcaniclastic sediments derived from the eastern Australian continental margin (Zirakparvar et al., 2013), and the metamorphic rocks of the Lousiades have been interpreted to represent mid-Cretaceous sediments scraped off of a subducting plate and incorporated into an accretionary wedge in the fore arc of a southward-migrating Miocene volcanic arc over northward dipping subduction. Woodlark Island may represent a remnant of the volcanic back-arc associated with this system (Webb et al., 2014). These scenarios would imply that emergent land masses may have been present in the Louisiades sector since at least the early Miocene.

### Woodlark island

This island is also currently *>* 200km from the Papuan mainland, lying in an isolated position to the northeast, and also contains a surprising number of Asterophryinae species, all with sister taxa on the EPCT (Fig. 4). Pigram and Davies (1987) treated Woodlark as lying on a separate terrane from the Louisiades, but noted that it had a basement of Eocene age, overlain by Oligocene limestones. Webb et al. (2014) suggested that Woodlark Island might represent a remnant of the volcanic back-arc associated with the Miocene arc that formed the Louisiades, which is consistent with its basement of pre-Miocene basalts reported by Ashley and Flood (1981), indicating that Woodlark and the Louisiades may have had a linked tectonic history. One hypothesis is that Woodlark was adjacent to the Louisiade Archipelago (Pigram and Davies, 1987; Baldwin et al., 2012), prior to the opening up of the Woodlark Basin which formed by rifting that initiated in the Late Miocene and continues to the present time (Webb et al., 2014). This rifting displaced the submarine ridge that Woodlark sits on to the northeast, pushing it away from its previous proximity to the Louisiades. This geological model is consistent with the large number of independent frog dispersals to Rossel island (currently the most easterly of the Louisiades), Sudest and Woodlark, between *∼*12–8 Ma, consistent with the hypothesized timing of fragmentation among these three islands. Thus the initial proximity of Woodlark Island to the Louisiades provided opportunities for dispersal and differentiation, which tapered off as the oceanic gap widened. Misima, which is the most westerly of the Louisiades and nearest to New Guinea, has a later history of dispersal from the EPCT, starting around *∼*6 Ma, consistent with a more recent separation.

### The D’Entrecasteaux archipelago

The D’Entrecasteaux group contains an exceptional diversity of asterophryne species on Normanby island, with more modest assemblages on Fergusson and fewer yet on Goodenough, which is a bit more xeric and less suitable for frogs. Asterophryinae dispersal to the D’Entrecasteaux Islands began *∼*8–5 Ma and continues to the present. These islands are an emergent metamorphic core complex, consisting of gneiss and amphibolite domes with granodiorite cores (Pigram and Davies, 1987), and with the potential exception of part of Normanby, were never connected to the mainland. They are not volcanic features, but instead seem to represent incipiently subducted continental crust that has rebounded back up through the zone of weakness at the west end of the Woodlark Rift (Baldwin et al., 2008; Wallace et al., 2014). They are also relatively new islands, of Pliocene age or later (Abers et al., 2002), and are still rising and enlarging. So unlike the Louisiades, which are old islands that are becoming progressively smaller, the D’Entrecasteaux group consists of young islands that are becoming progressively larger. As such, they are a very recent colonization target that has required overwater dispersal to reach.

Goodenough and Fergusson islands are internally cohesive emergent metamorphic units, while Normanby Island has a more complicated and potentially composite history. The island is divided by a right-lateral strike-slip fault (Wallace et al., 2014). The narrower western section of Normanby, to the north of this fault, is of similar composition and origin to Goodenough and Fergusson islands, while the larger, furrowed eastern section is a separate lithological unit that is more allied to parts of the Owen Stanley uplift in the central Papuan Peninsula (Baldwin et al., 2012). Thus, the section of Normanby where our samples were collected, with a surprising species richness (24 candidate species), may be of separate origin more closely linked to the EPCT possibly representing a younger version of the situation in the Louisiades. The different tectonic histories of Normanby versus Fergusson and Goodenough help to explain the species diversity disparities, while providing evidence that overwater dispersal can occur when short distances are involved.

### New Britain island

Currently there are only two species of Asterophryinae known to be present on New Britain, both of which dispersed there within the past 3 Ma, and have sister taxa on both the Accreted Terranes and the EPCT. Considering that New Britain is on a collision course with the Papuan mainland, and currently lies as close as it ever has to New Guinea (Kroenke et al., 1984; Pigram and Davies, 1987; Polhemus and Polhemus, 1998; Quarles van Ufford and Cloos, 2005; Baldwin et al., 2012; Davies, 2012), overwater dispersal is the only mechanism available to frogs, and the low number of asterophryne species despite its large land area makes sense in this context.

### South East Asia

The historical genus *Oreophryne* was recently split into two distinct and unrelated genera *Oreophryne A* which is the oldest genus of Asterophryinae, and a younger, unrelated clade *Oreophryne B* (Hill et al., In Review). We have four *Oreophryne* samples from this region, two species in *Oreophryne A* from the Philippines which are sister to species on the Accreted Terranes, and two species in *Oreophryne B* from Sulawesi which are sister to species on the EPCT. Dispersal to Sulawesi could be explained by westward-motion along the Pacific plate boundary, where accommodation of convergence has sheared fragments from northern New Guinea and sent them westward towards the Sulawesi region (Hamilton, 1979; Polhemus, 2007) along a series of left-lateral fault zones. These cases suggest an out-of-PNG to the Sunda Arc dispersal route hypothesis. Recently, several additional genera of Asterophryinae from South East Asia have been described: *Siamophryne, Gastrophrynoides*, and *Vietnamophryne* and were found to be basal sisters of the Papuan Asterophryinae (Poyarkov et al., 2018; Suwannapoom et al., 2018). This was interpreted as evidence for a “down-from-Southeast-Asia” pattern of dispersal (Tyler, 1979). More sampling of species as well as inclusion of nuclear genes in phylogenetic analyses is required to resolve the structure of the South East Asian clades in relation to the Papuan clades and to determine whether such faunal disjunction might be linked to much earlier tectonic events.

### A Comment About Model Based Approaches to Biogeography

We employed a phylogenetic model-based approach to explore hypotheses about historical biogeography. That is, we constructed a series of models that approximate our hypothetical phenomena (i.e., distance, various geological scenarios), and allowed them to compete for the best explanation of the data. Luckily, despite the complexity of hypotheses involving partial areas, it appears that our model fits are additive such that for example – the addition of Woodlark to the Louisiades produces the same improvement of fit regardless of the assumptions of other parts of the model. We caution that this need not be the case, and care must be taken in making such assessments. Nevertheless, the gains for biology in rigorous interrogation of models are tremendous. By comparing models head to head, we could clearly identify which ideas were supported by data, and which were not.

It has been noted by Ree and Sanmart໭ n (2018) that many studies do not exploit the power of model specification flexiblity afforded by the DEC class models. Rather, most studies limit the comparison to major classes of models (i.e., DEC, DEC+J, DIVA), but do not vary the model to approximate the actual phenomena of interest, leaving conclusions to interpretation of the ancestral range reconstructions. Interpretation of model results is, of course, very important, we agree that it is a missed opportunity to use the power of statistical model-comparison approaches to drill down into the exploration of the data. We can use a hypothesis-driven phylogenetic approach to differentiate between customized hypotheses to test specific biological or geological phenomena.

## Conclusions

We find using an explicit model-based approach clear signals in the data with very strong support for each of these model additions: the pairwise connections of the mainland, the affiliation of the Louisiades+Woodlark with the EPCT, as well as the D’Entrecasteux. Together with the interpretation of the ancestral range reconstructions and their coincidence in timing with geological events, they support the “Slow and Steady” geological hypothesis for the formation of the region (Pigram and Davies, 1987; Davies et al., 1996; Polhemus and Polhemus, 1998; Quarles van Ufford and Cloos, 2005; Davies, 2012), with the EPCT as the center of origin and source for ongoing diversification. Within this geological context, we find no signal for elevational specialization, but we do find support for the role of isolation once geological history is taken into account. In a group with impossibly complex distribution yet with poor dispersal ability on islands with no current connection to mainland, we need not accept wild dispersal scenarios. We can rigorously test geological ideas to find a strongly supported biogeographical model involving good old fashioned land connections (in multiple varieties of joining and fragmenting) to explain the convoluted dispersal pathways and exhuberant diversity of the Asterophryinae.

Several authors have noted that island biogeography theory is less predictive over evolutionary timescales. Given sufficiently long timescales, other processes may dominate over an equilibrium between immigration and extinction, such as: the accumulation of in-situ radiation (Cowie, 1995; Heaney, 2000), evolutionary specialization and changes in food web structure (Borges and Brown, 2008), and geological changes in land area (Whittaker et al., 2008). A major difference between our study and those listed above is that they use (non-phylogenetic) multiple regression-based approaches to analyze correlations between explanatory variables and species richness. While ahistorical approaches have the advantage of inclusion of a broader diversity of taxa to address a general phenomenon, a phylogenetic approach offers greater precision in exploring both time separating the lineages and where they occur in space. We can then explore a greater variety of hypotheses involving complex spatio-temporal processes. Ultimately, we do find support for some of the predictions of the theory of island biogeography over evolutionary time scales if the geological history of plate tectonics is considered. The earth’s surface is not fixed but itself evolves. Not only can land area change over time, sometimes increasing by volcanism or shrinking through erosion, but the distances between land masses can evolve as well, changing the degree of isolation through time and providing another axis upon which geological evolution can alter the dynamics of biotic evolution.

## Acknowledgements

This study was made possible by a grant from the National Science Foundation awarded to MB (DEB-1145733). Julio Rivera, Jeff Scales, Nalani Kito-Ho, Niegel Rozet, Jeff Higa and Raine Higa provided generous assistance in the field. We are grateful to the field assistance from many local residents: at Maru Ruama, Mt. Geregu: Peter Joseph, Dambio Moi, Walter Moi, David Peter, and Laiwoi Yoiini; at Cliffside Camp, Kamiali: Maties Dagam, David Enoch, Lenny Keputong, and Marcus Symon; at Normanby Island: Clement Bobby, Fred Francisco, Kenny Lakson, Waiyaki Nemani, James Waraia, and Roland Waraia, and Misima Island: Simon Emidi and Kelly Nabwakulea. We thank Normanby Mining PNG, LTD for land access and accommodation. We thank Georgia Kaipu of the PNG National Research Institute) and the late Barnabus Wilmont of the PNG Department of Environment and Conservation for assistance with research permits and necessary visas, and Andrew Moutu, the Director of the PNG National Museum and Art Gallery, for use of facilities.

## Supplementary Materials

**Fig. S1.**
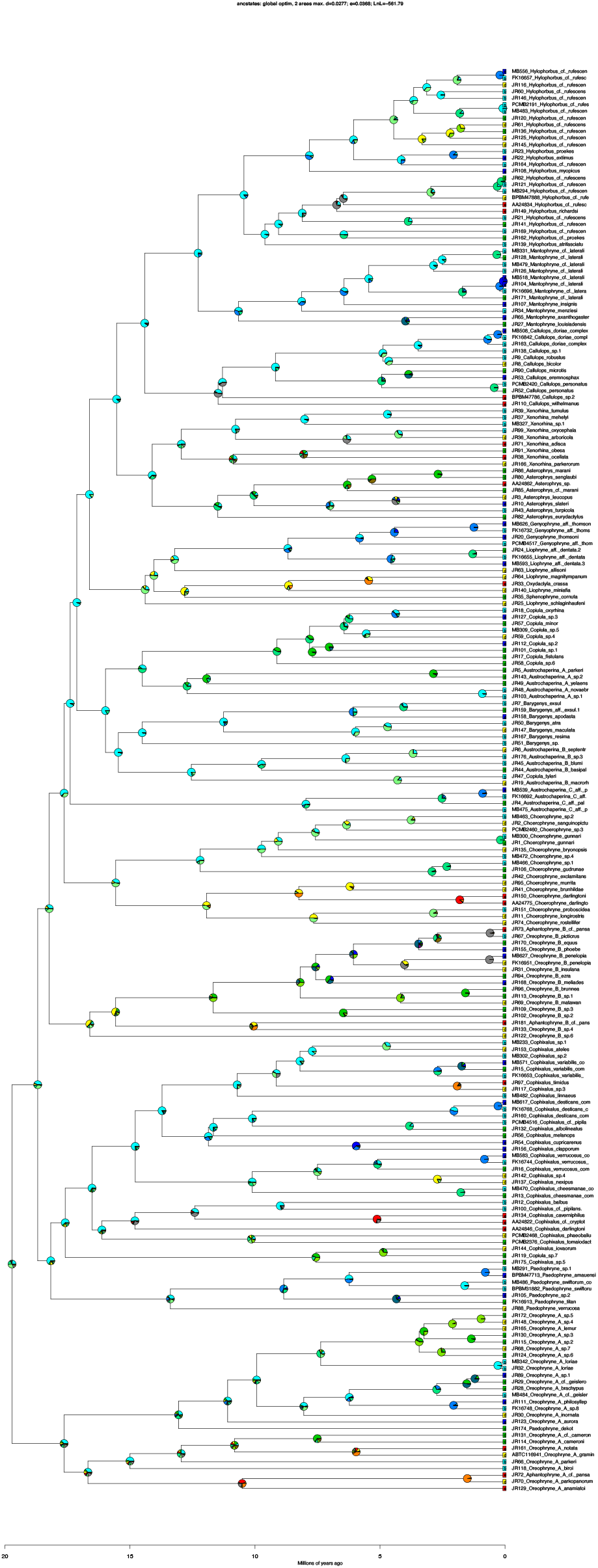
Supplementary Figure 1: Elevational range evolution in Asterophryinae. Ancestral elevational states were reconstructed using DEC models.

**Fig. S2.**
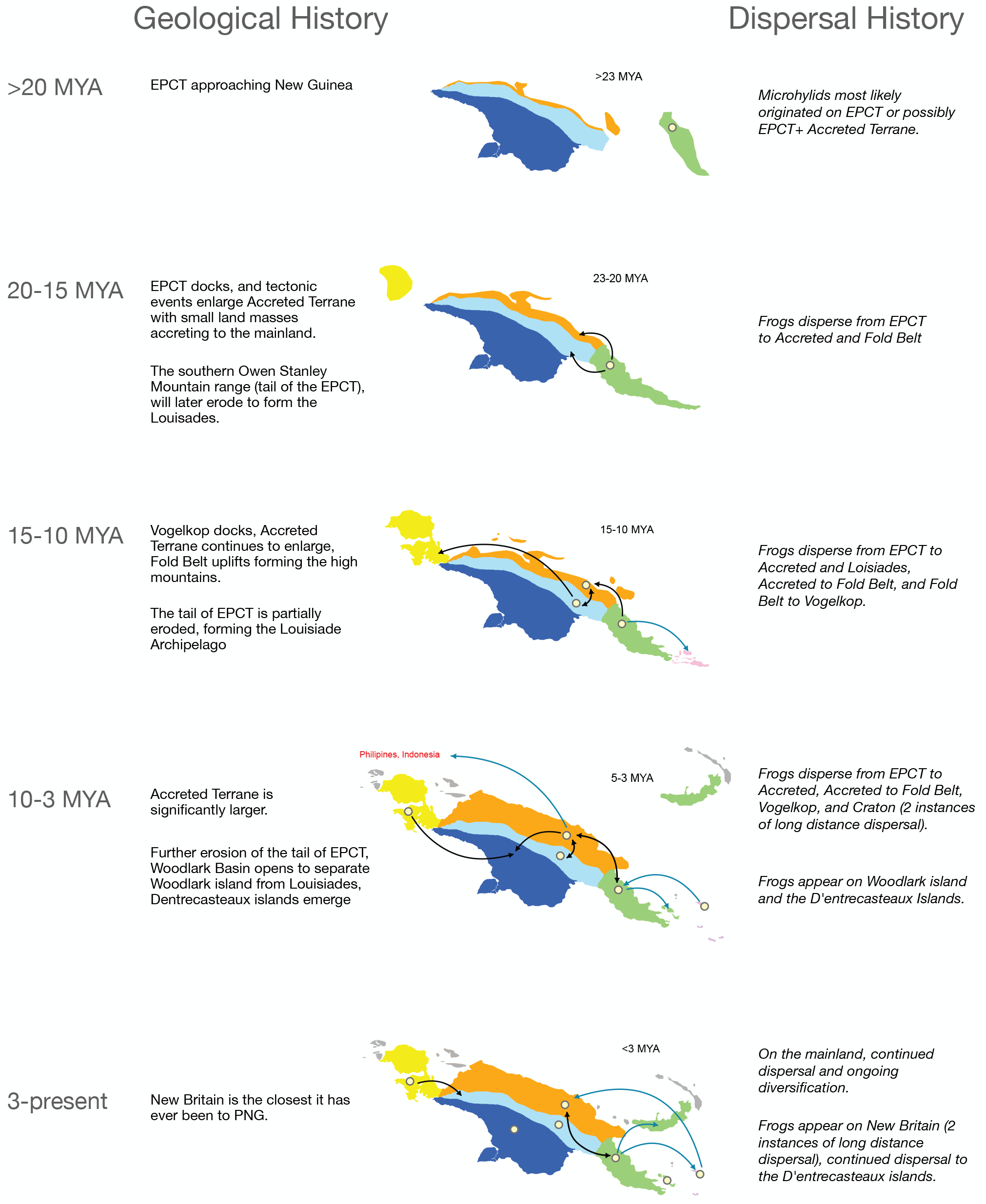
Narrative of the geological history of the mainland over the last 20MY (Davies, 2012) and its offshore islands (Pigram and Davies, 1987), along with dispersal pattern results from this study of Asterophryinae through time.

